# Differential and lasting gene expression changes in circulating CD8 T cells in chronic HCV infection with cirrhosis and related insights on the role of Hedgehog signaling

**DOI:** 10.1101/2023.09.20.557725

**Authors:** Jiafeng Li, Agatha Vranjkovic, Daniel Read, Sean P Delaney, William L Stanford, Curtis L Cooper, Angela M Crawley

## Abstract

The impact of chronic hepatic infection on antigen non-specific immune cells in circulation is not well understood and may influence long term health. We reported lasting global hyperfunction of circulating CD8 T cells in HCV-infected individuals with cirrhosis. Whether gene expression patterns in bulk CD8 T cells are associated with the severity of liver fibrosis in HCV infection is not known. RNA sequencing of blood CD8 T cells from treatment-naïve, HCV-infected individuals with minimal (Metavir F0-1 ≤ 7.0 kPa) or advanced fibrosis or cirrhosis (F4 ≥ 12.5 kPa), before and after direct-acting antiviral therapy, was performed. Principal component analyses determined robust differences in over 350 genes expressed by CD8 T cells from HCV-infected individuals with minimal or advanced fibrosis and data suggests this remains relatively stable after viral clearance. Gene ontology analyses identified disaggregated gene expression related to cellular metabolism, including upregulated phospholipase, phosphatidyl-choline/inositol activity and second-messenger-mediated signaling, while genes in pathways associated with nuclear processes, RNA transport and cytoskeletal dynamics were reduced. Gene Set Enrichment Analysis identified decreased expression of genes regulated by the cMyc and E2f transcription factors in cirrhotics, compared to the minimal fibrosis group, as well as reduced expression of genes linked to oxidative phosphorylation, mTOR signaling, and more. Upregulated gene sets in cirrhotics included IFN-α, -γ, TGF-β response genes, apoptosis and apical surface pathways, among others. The hedgehog (Hh) signaling pathway was the top featured gene set upregulated in cirrhotics. Inhibition of Hh signaling with cyclopamine ablated CD8 T cell IFN-γ production, suggesting its involvement in hyperfunction. This is the first analysis of bulk CD8 T cell gene expression profiles in HCV infection in the context of liver fibrosis severity, and suggests cirrhosis significantly reprograms the CD8 T cell pool. The novel finding of increased Hh signaling in cirrhosis may contribute to generalized CD8 T cell hyperfunction observed in chronic HCV infection. Understanding the lasting nature of immune cell dysfunction may help mitigate remaining clinical challenges after HCV clearance and more generally, improve long term outcomes for individuals with severe liver disease.

## INTRODUCTION

Available statistics on the prevalence of advanced stage liver disease and cirrhosis suggest significant disease burden worldwide, for which hepatic viral infections and metabolic dysfunction-associated steatotic liver disease (MASLD, formally named nonalcoholic fatty liver disease) are the leading causes. As antiviral treatments for hepatic viral infections have emerged to either eliminate or control viremia, as with hepatitis C (HCV) and B (HBV), respectively, remaining challenges for affected individuals include the long-term outcomes of cirrhosis. Once liver disease progresses to advanced liver fibrosis or cirrhosis, there is an increased risk for progression to end-stage liver disease, portal hypertension, esophageal varices, susceptibility to infections, and hepatocellular carcinoma (HCC). While several studies have suggested that direct-acting antiviral (DAA) therapies have reduced the risk for HCC in HCV-infected individuals^1–4^, there still remains appreciable risks of HCC development following DAA therapy, as well as HCC recurrence following anticancer treatment^5^. In addition to the associated metabolic disease in cirrhosis, a variety of immune dysfunctions emerge as liver disease progresses, affecting both innate and adaptive immune responses^6^. While innate immune dysfunctions have been well described in cirrhosis^7–9^, the contribution of adaptive immune defects to the health outcomes of cirrhosis has not been fully examined.

Chronic HCV infection disrupts immune functions, affecting many innate and adaptive immune cells, including cytotoxic CD8 T cells^10–15^. In the acute stage of HCV infection, weak and transient responses of HCV-specific CD8 T cells predict chronicity^16,17^. In chronic infection, detection of impaired HCV-specific CD8 T cells prior to IFN-α + ribavirin antiviral therapy predicts return to chronic infection if reinfected^18^, as these cells remain dysfunctional^12,19^, suggesting irreversible damage to immune cells. While DAA therapies have achieved spectacular results for viral clearance, it is not clear if viral cure parallels with restored immune functions given contrasting reports in the literature^20,21^. In addition, HCV infection has a more extensive, antigen agnostic effect on CD8 T cells, as markers of exhaustion are widely observed on bulk CD8 T cells in the blood, spleen and liver^22–25^. A study has observed exaggerated proliferation, cytokine secretion and degranulation by *in vitro-*stimulated cytomegalovirus or Epstein-Barr virus (CMV/EBV)-specific CD8 T cells in HCV-infected individuals and this was retained after DAA therapy^26^. However, the effects of liver fibrosis severity on the acquisition and possible long-term retention of T cell dysfunction have not been determined. Attempts to restore normal function *in vitro* in isolated HCV/CMV/EBV-specific CD8 T cells from HCV-infected individuals with advanced liver fibrosis have failed^27^. This suggests that the immune system is profoundly affected in HCV infection according to the degree of liver fibrosis.

We and others have observed significant impairment of the entire CD8 T cell compartment in the blood and liver in HCV infection^22–25^, wherein we specifically associated decreased CD8 T cell survival with advanced liver fibrosis^28^. We then showed for the first time an overactive bulk CD8 T cell function profile in HCV-infected individuals with cirrhosis compared to those with minimal fibrosis^29^. This was done alongside our complimentary clinical study in which a cohort of DAA-treated HCV^+^ patients with cirrhosis achieved a sustained virological response, SVR (i.e. undetectable HCV RNA by 12 weeks after treatment cessation) yet failed to reverse liver fibrosis by 24-weeks after viral clearance^30^. We hypothesized that liver fibrosis severity is strongly associated with immune dysfunction. Consistent with this theory, we found that bulk CD8 T cell responses were not restored to physiological levels after DAA therapy in cirrhotic individuals^29^.

Characterizing the underlying mechanisms of bulk CD8 T cell dysfunction in advanced liver fibrosis is of significant clinical importance given the role of CD8 T cells in response to infection and cancer surveillance. In this study, we identify several candidate genes and pathways that may contribute to this hyperfunction and inform future mechanistic investigations, namely Hedgehog (Hh) signaling. Hh signaling is widely recognized as an important component of embryonic development and tissue regeneration^31,32^. The signaling cascade is initiated by extracellular Hh ligands (homologs Sonic, Desert and Indian, respectively Shh, Dhh, and Ihh) binding to receptor Patched-1 (Ptch-1) or Patched-2 (Ptch-2),and results in, via interaction with Smoothened (Smo), downstream activation of Gli transcription factors (Gli1, Gli2, and Gli3). Non-canonical Gli-independent Hh signaling also plays a prominent role in Ca^2+^ signaling and cytoskeletal rearrangement^33^. More recently, it has been shown that exposure to Shh ligand produced by thymic epithelial cells in the adult is essential to drive differentiation and proliferation of thymocytes during transition from the double-negative 1 to 2 stage of development^34^. Hh signaling was also found to support γδ T cell maturation^35^. In mice, Hh signaling was also shown to be involved in the formation of the immunological synapse of CD8 T cells^36^. This study is thus complemented with data showing a strong dependence on Hh signaling in CD8 T cell function and its contribution to immune cell hyperfunction in chronic HCV with advanced liver fibrosis.

## RESULTS

### CD8 T cell gene expression profiles between advanced and minimal fibrosis in chronic HCV^+^ individuals differ greatly and are independent of HCV clearance

An analysis of HCV-infected subjects based on liver fibrosis severity before and after DAA therapy has been rarely conducted in the study of bulk CD8 T cells. Our previous work demonstrated of CD8 T cell hyperfunction in blood cells from HCV-infected individuals with advanced liver fibrosis prior to DAA therapy, compared to those with minimal fibrosis^29^. The gene expression profiles of 8 individuals (Table 1) infected with HCV, and naïve to treatment, were examined, of whom 4 were identified as having minimal liver fibrosis (F0-1, ≤ 7.0 kPa) before the initiation of DAA therapy and 4 had developed advanced fibrosis/cirrhosis (F4, ≥ 12.5 kPa) before treatment. All treated individuals cleared the virus (i.e., achieved sustained virological response, SVR).

**Table 1:** Summary of demographics and clinical information of study groups.

| Parameter | Samples used in RNA-sequencing |  | Samples used in PCR and T cell functional analyses |  |  |
| --- | --- | --- | --- | --- | --- |
|  | HCV<sup>+</sup> AF (F3-F4)<sup>a</sup> | HCV<sup>+</sup> MF(F0-1)<sup>a</sup> | HCV<sup>+</sup> AF (F3-F4)<sup>a</sup> | HCV<sup>+</sup> MF (F0-1)<sup>a</sup> | Uninfected |
| Sample size | 4 | 4 | 4 | 4 | 4 |
| Sex | 3 M, 1 F | 1 M, 3 F | 3 M, 1 F | 3 M, 1 F | 2 M, 2 F |
| Mean age ± SD (Range) | 58.5 ± 12.4 (46–75) | 48.5 ± 5.0 (41–51) | 53.8 ± 7.1 (46–62) | 52.5 ± 15.2 (33–65) | 47.5 ± 10.6 (35–59) |
| Ethnicity | Caucasian (3),<br>S-E Asian (1) | Caucasian (4) | First Nations (2),<br>Caucasian (1),<br>Unknown (1) | Caucasian (3),<br>Unknown (1) | Caucasian (2),<br>South-American (1),<br>Unknown (1) |
| Mean fibrosis score<sup>b</sup> (kPa) ± SD | 23.9 ± 13.6 | 3.9 ± 1.7 | 16.2 ± 3.4 | 5.3 ± 1.1 | n/a |

<sup>a</sup>AF = Advanced fibrosis, Metavir F3-F4 ≥ 12.5 kPa<sup>b</sup>; MF = Minimal fibrosis, Metavir F0-1 ≤ 7.0 kPa<sup>b</sup>.
<sup>b</sup>Liver stiffness in kilopascal (kPa) measured by Fibroscan transient elastography.

We isolated bulk CD8 T cells from PBMC samples collected from these study groups pre-DAA and 12 weeks post-SVR, followed by 16h stimulation with 5μg/ml phytohaemagglutinin (PHA) and RNA-sequencing. In total, 24,058 detectable genes were retained for analysis out of a database of 58,294. Hierarchical clustering based on normalized gene expression counts across all transcripts identified a clear clustering between samples from individuals with advanced fibrosis compared to minimal fibrosis (Fig. 1A). A principal component analysis (PCA) applied to the top 500 most variable genes in this dataset revealed clear differences in gene expression in patients with advanced fibrosis before treatment compared to that of individuals with minimal fibrosis (Fig. 1B). PCA broadly separated the minimal fibrosis patients (patients #116, 117 and 137) from the advanced fibrosis/cirrhotic patients (patients #133, 136 and 171), with one exception (patient #130). Where applicable, the CD8 T cell gene expression profiles of treatment-paired patient samples (pre-vs. post-DAA) remained constant in the three individuals with minimal liver fibrosis (patients #116, 117 and 137) as well as the single DAA-treated individual with high fibrosis analyzed in this study (patient #133). Fold-change differences based on fibrosis severity before treatment moderately correlated with severity-based differences after treatment (Fig. 1C), indicating no significant change between gene expression profiles before and after DAA treatment.

**Figure 1:**
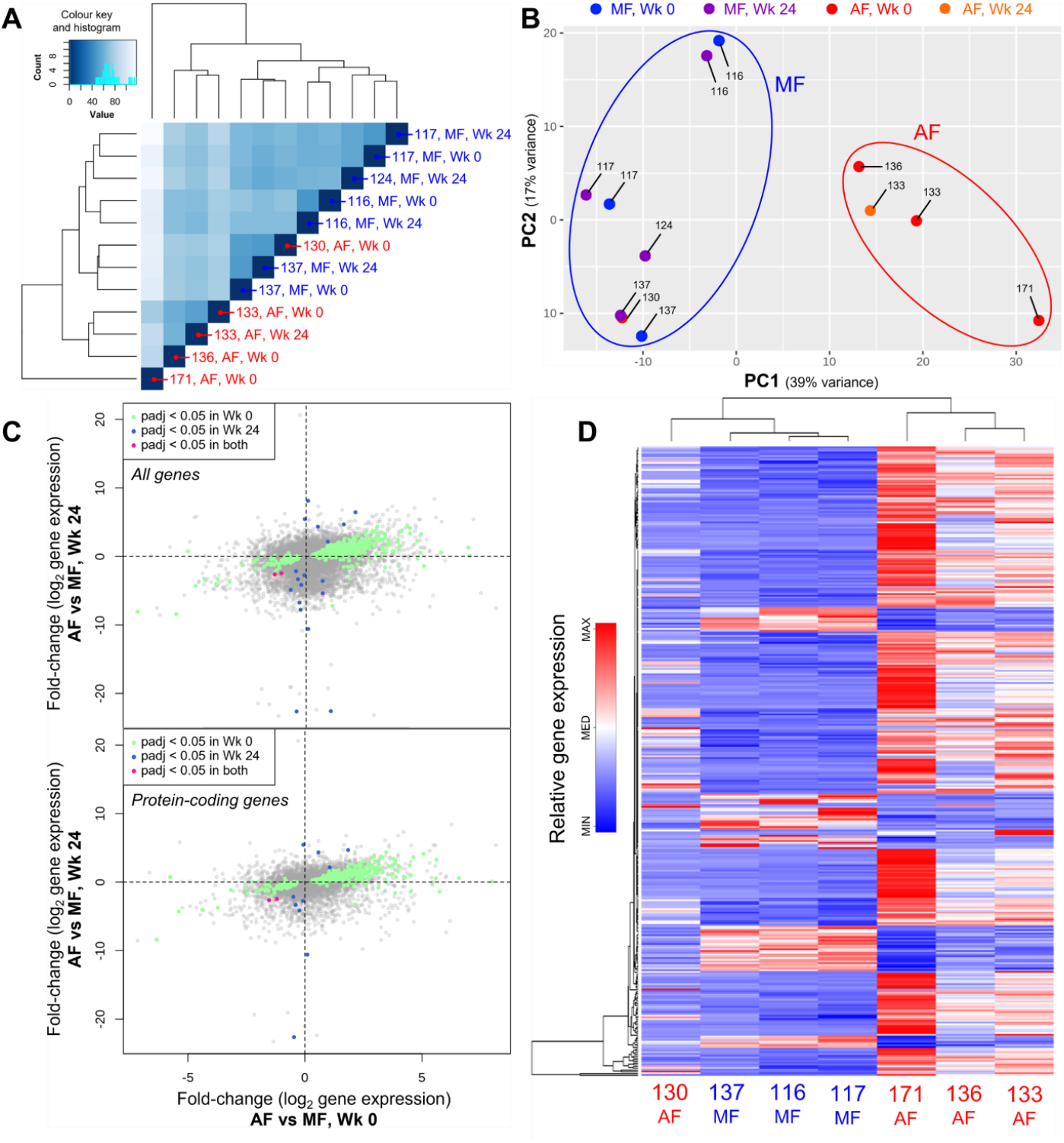
Gene expression profiles of CD8+ T cells in chronic HCV patients with advanced fibrosis (AF) differ greatly from minimal fibrosis (MF) patients. Isolated CD8 T cells were stimulated with 5 μg/ml PHA for 16h prior to RNA-sequencing. A) Clustering of samples based on all RNA-seq transcripts results in a clear separation between AF and MF regardless of treatment stage. B) PCA applied to the top-500 most variable genes separates AF and MF patients in distinct clusters, irrespective of treatment stage. C) Plotting gene expression fold-change differences before DAA therapy (Wk 0) compared to 12 weeks post-DAA therapy (Wk 24) shows an association across all genes significantly differentially expressed (top), as well as well as differentially expressed protein-coding genes (bottom). D) Heatmap clustering of all 362 genes significantly different between AF and MF patients show clear distinct gene expression profile patterns between these study groups.

In pre-treatment individuals with advanced fibrosis compared to minimal fibrosis, 362 genes were significantly differentially expressed (p-adj<0.05), of which 288 genes were upregulated and 74 were downregulated. Complete-linkage clustering of these genes separated CD8 T cell gene expression profiles of advanced fibrosis individuals (patients 133, 136, 171) from minimal fibrosis (patients 116, 117, 124, 137), with patient 130 as the exception (Fig. 1D), consistent with PCA. Comparison of gene expression profiles between these study groups at 24 weeks into DAA treatment identified only 20 significant genes (6 upregulated, 14 downregulated). Clustering of these genes also separated individuals with advanced fibrosis separately from minimal, although less noticeably (data not shown).

Taken together, this suggests that CD8 T cell gene expression is greatly affected by the severity of liver fibrosis in chronic HCV infection, and that this difference tends to remain long after DAA-mediated viral clearance. To obtain high quality, generalizable results regarding liver fibrosis severity effects on bulk CD8 T cells during chronic HCV, analyses on adequately grouped and sufficiently large datasets is vital. Therefore, the remainder of this investigation focuses on untreated HCV-infected individuals evaluated based on liver fibrosis severity (cirrhosis vs. minimal).

### Differential gene expression in bulk CD8 T cells in HCV infection with advanced/minimal liver fibrosis is associated with cellular metabolism and cell structure and motility

To determine which cellular functions are modulated by these gene expression patterns, we performed a functional enrichment analysis of the lists of differentially expressed genes in HCV-infected individuals with cirrhosis of minimal fibrosis before DAA treatment. The enrichment analysis searched databases of functional categories and highlighted gene sets that may be statistically over-represented in the dataset. The adjusted *p*-value cutoff of 0.1 was used to identify enriched groups. Each term in these analyses contains a set of genes with correlated expression patterns, annotated by biological function.

An analysis for Gene Ontology (GO) Molecular Function (MF) and Biological Processes (BP) classifications with ≥ 3 enriched genes per term was performed. GO MF terms associated with upregulated genes in the cirrhosis group compared to the minimal fibrosis group include phospholipase activity (*p* = 0.003), lipase activity (*p* = 0.011), and phospholipase C activity (*p* = 0.053), amongst others (Fig.2A). GO BP terms associated with upregulated genes include second-messenger-mediated signaling (*p* = 0.026), and regulation of leukocyte migration (*p* = 0.040), amongst others (Fig.2A). An enrichment analysis in Kyoto Encyclopedia of Genes and Genomes (KEGG) pathways and modules was also performed. Notable KEGG hits enriched for upregulated gens in cirrhosis include NK cell-mediated cytotoxicity (*p* = 0.043), and phospholipase D signaling (*p* = 0.043), amongst others (Fig. 2A). Following GO analyses, gene set enrichment analysis (GSEA) was conducted on significantly differentially expressed genes in cirrhosis. In total, 10 GSEA upregulated gene sets were enriched (FDR q ≤ 0.1) in CD8 T cells from cirrhotic individuals compared to minimal fibrosis, including Hedgehog signaling (14 genes, including PTCH1 and GLI1), IFN-α and -γ responses (31 and 59 genes respectively), and apoptosis (51 genes), amongst others (Fig.2B-C). In genes downregulated in the cirrhosis group, no GO MF terms nor KEGG pathways or modules were enriched, while GO BP terms enriched include nuclear division (p = 0.013), actin nucleation (p = 0.017), as well as RNA transport (p = 0.032) and localization (p= 0.056), amongst others (Fig.2D). GSEA hits in downregulated genes include Myc and E2F targets (54 and 130 genes respectively), oxidative phosphorylation (122 genes), G2/M checkpoint (112 genes), mTORC1 signaling (93 genes), and DNA repair (68 genes), amongst others (Fig.2E-F). Overall, these analyses indicate that the gene expression differences in CD8 T cells from chronic HCV patients with cirrhosis spread across many vital signaling pathways and cellular processes. These differences hinge mostly upon functions associated with cellular metabolism and cell growth, T cell activation and inflammatory responses, RNA transport, as well as cytoskeletal control and cellular migration.

**Figure 2:**
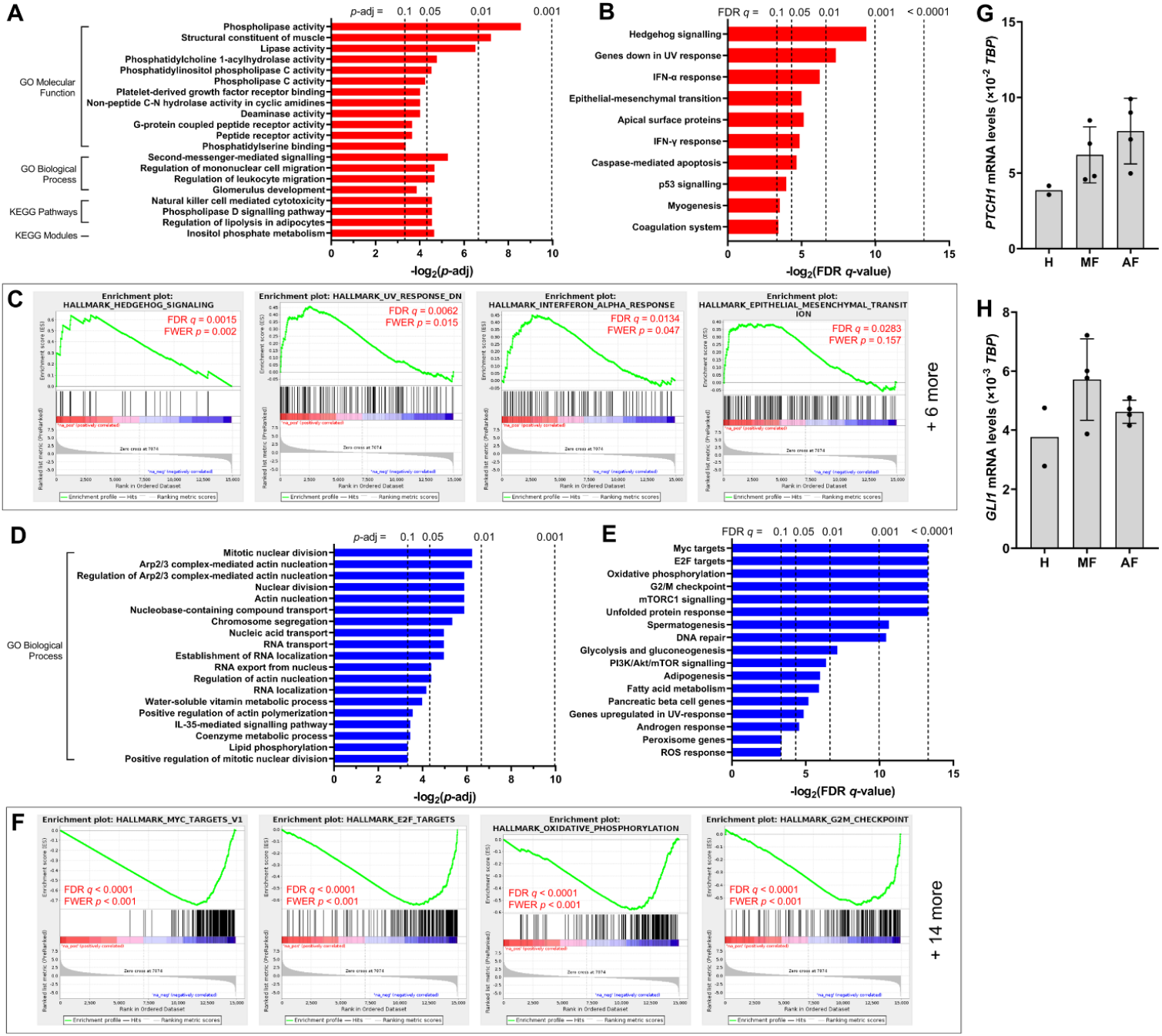
Functional enrichment analysis of RNA-seq data indicate changes in pathways related to CD8 T cell responses and metabolism in pre-DAA HCV patients with advanced fibrosis (AF) compared to minimal fibrosis (MF). A)GO analysis of upregulated genes in CD8 T cells from AF patients compared to MF identifies multiple classifications related to metabolic regulation. B) GSEA enriched 10 gene sets upregulated in AF, notably hedgehog (Hh) signalling and inflammatory responses. C) GSEA enrichment plots of the top four hits in upregulated gene sets in AF compared to MF. D) GO analysis of downregulated genes in AF identifies multiple classifications related to cytoskeletal regulation and nucleic acid transport regulation. E) GSEA enriched 18 gene sets (including two versions of Myc targets, both FDR q<0.0001) downregulated in AF, many of which relate to metabolic regulation. F) GSEA enrichment plots of the top four hits in downregulated gene sets in AF compared to MF. G-H) Relative mRNA expression of Hh signalling components PTCH1 and GLI1 in isolated CD8 T cells from AF or MF patients, compared to cells from healthy individuals (H), assessed by qPCR after 16h of stimulation using anti-CD3/CD28 antibodies.

### Inhibition of canonical Hedgehog signaling ablates CD8 T cell hyperfunction in advanced fibrosis

Given the importance of Hh signaling in mediating T cell development and cytotoxic functions^34,37,36^, we confirmed differential RNA expression in CD8 T cells in disparate liver fibrosis contexts in HCV infection by qPCR analysis of PTCH1 and GLI1 mRNA. In isolated CD8 T cells stimulated with anti-CD3/CD28 antibodies for 16h, PTCH1 and GLI1 mRNA expression was elevated in HCV^+^ individuals relative to uninfected healthy controls, and PTCH1 mRNA expression levels associated with fibrosis severity (Fig. 2G-H).

We next examined whether Hh signaling contributes to IFN-γ and perforin expression in CD8 T cells. Circulating CD8 T cells isolated from HCV patients and healthy individuals (Table 1) were stimulated for 48h with anti-CD3 and anti-CD28 antibodies, with or without cyclopamine, a chemical inhibitor of Smo and IFN-γ and perforin expression were measured by flow cytometry. Mirroring previously reported observations, CD8 T cells from cirrhotic individuals exhibit hyperfunction via increased IFN-γ and perforin expression across various cell subsets, notably in naïve cells and effector cells (Fig. 4). Cyclopamine treatment (50 μM) of CD8 T cells alongside anti-CD3/CD28 stimulation ablated IFN-γ expression in bulk CD8 T cells regardless of liver disease severity (Fig.3A-B). This dependence on Smo-mediated Hh signaling was also observed in naïve and effector cells in cirrhotics where hyperfunction was observed (Fig.3C-D). In naïve CD8 T cells, there was a trend between the degree of IFN-γ reduction and the degree of fibrosis severity (Fig.3C, ANOVA interaction p = 0.08). Perforin expression by CD8 T cells treated with cyclopamine was also ablated in bulk cells (Fig.4A-B), again occurring in naïve and effector cells (Fig.4C-D) where hyperfunction was observed and degree of reduction in both naïve (ANOVA interaction) and effector cells was also coupled to fibrosis severity (p = 0.09 and 0.03, ANOVA, respectively). Similar findings were observed with vismodegib, another chemical inhibitor of Smo (not shown). These results suggest that canonical Hh signaling plays an important role in IFN-γ and perforin expression during the CD8 T cell response and may play a role in bulk CD8 T cell hyperfunction in advanced fibrosis in HCV^+^ individuals.

**Figure 3.**
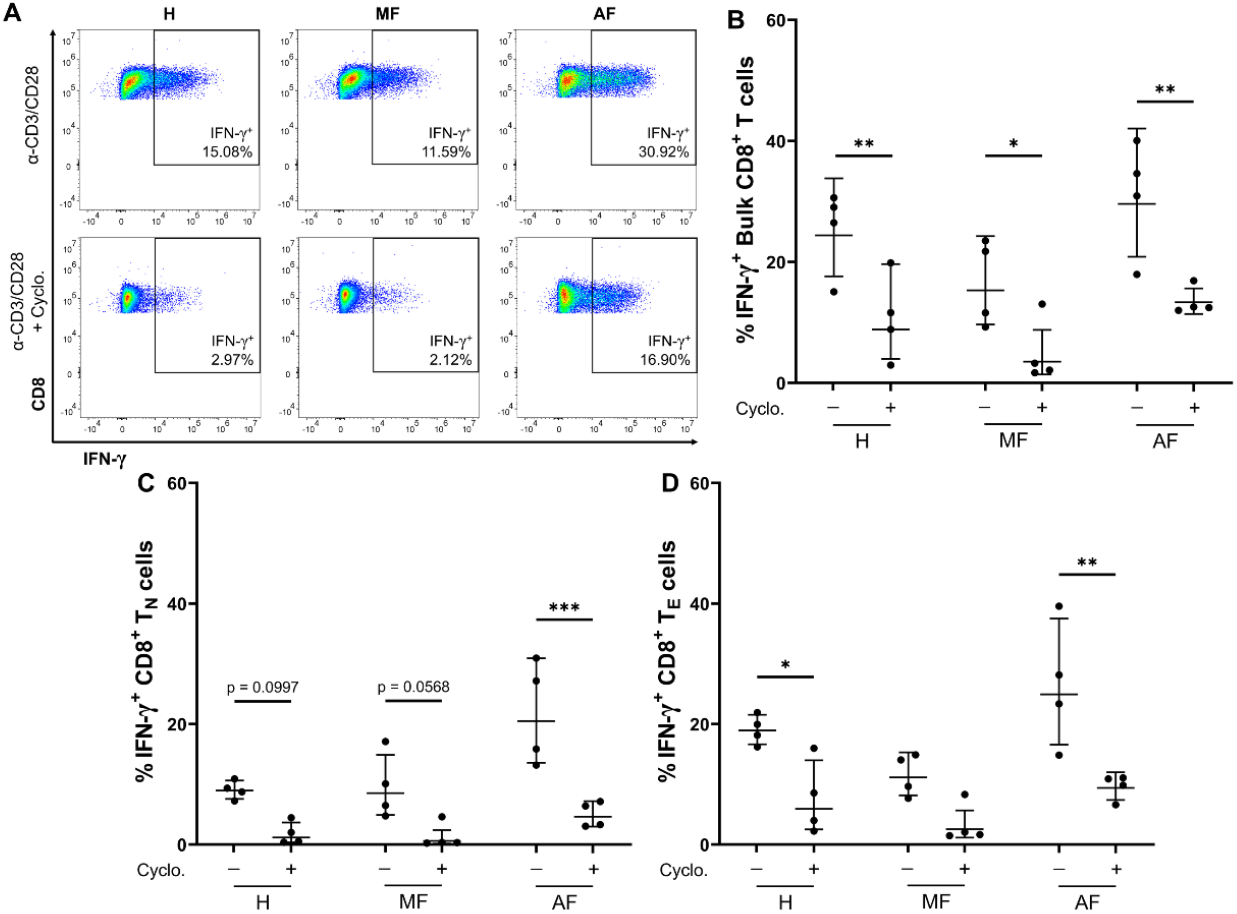
Inhibition of canonical Hedgehog signalling ablates IFN-γ expression in CD8 T cells, notably in naïve and effector CD8 T cells. A) Representative dot plots of IFN-γ expression in bulk CD8 T cells from HCV patients with advanced fibrosis (AF), minimal fibrosis (MF), or healthy individuals (H), stimulated for 48h anti-CD3/CD28 antibodies with or without 50 μM cyclopamine. B) Smo inhibition with cyclopamine during stimulation ablated IFN-γ expression in bulk CD8 T cells across all disease severities. C) Smo inhibition with cyclopamine during stimulation ablated IFN-γ expression in naïve CD8 T cells. The inhibition significance increases with hyperfunction in AF compared to MF or H (ANOVA interaction p=0.0833). D) Smo inhibition with cyclopamine during stimulation ablated IFN-γ expression in effector CD8 T cells. All multiple comparisons are analyzed by 2-way ANOVA with Šídák’s post-test, \**p*<0.05, \*\**p*<0.01, \*\*\**p*<0.001.

**Figure 4.**
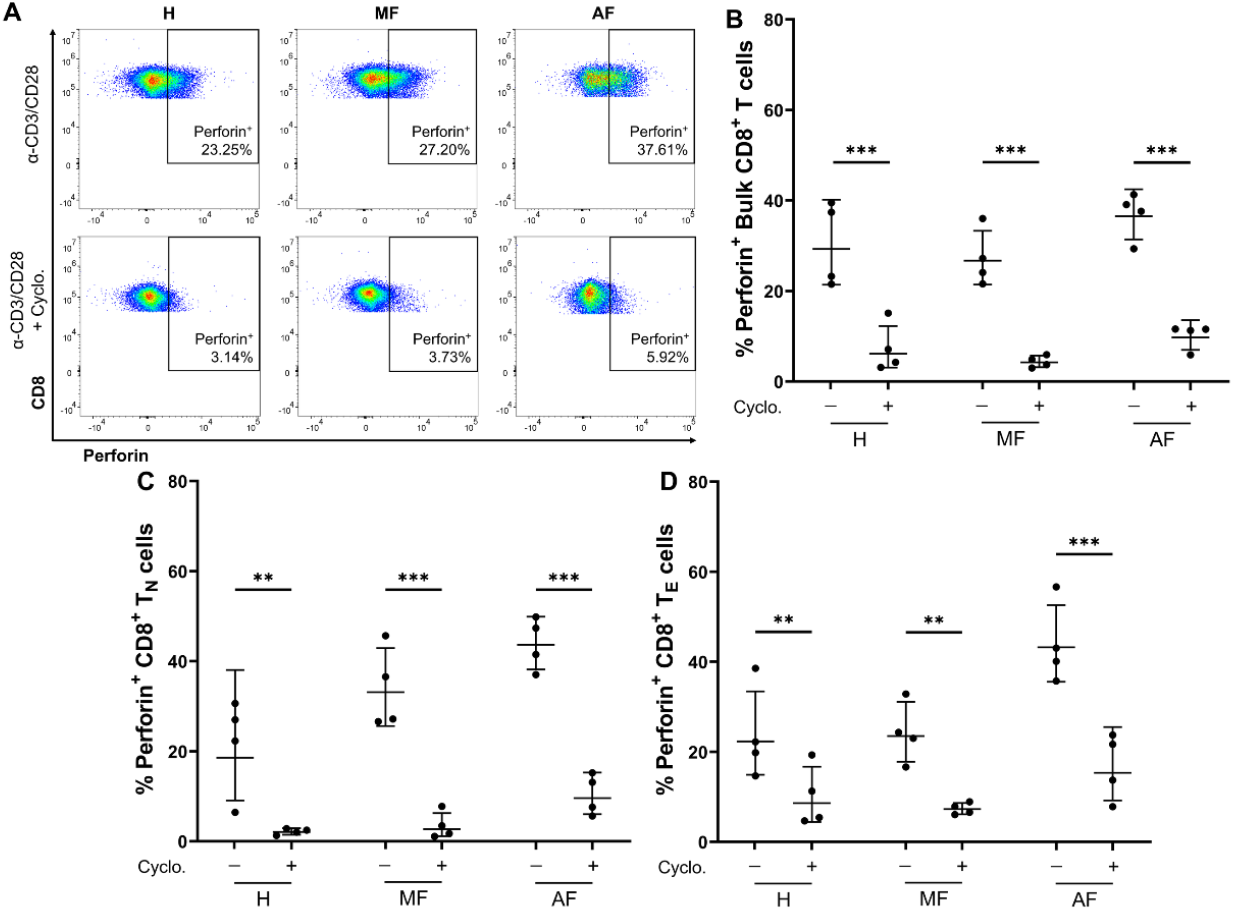
Inhibition of canonical Hedgehog signalling ablates perforin expression in CD8 T cells, notably in naïve and effector CD8 T cells. A) Representative dot plots of perforin expression in bulk CD8 T cells from HCV patients with advanced fibrosis (AF), minimal fibrosis (MF), or healthy individuals (H), stimulated for 48h anti-CD3/CD28 antibodies with or without 50 μM cyclopamine. B) Smo inhibition with cyclopamine during stimulation ablated perforin expression in bulk CD8 T cells across all disease severities. C) Smo inhibition with cyclopamine during stimulation ablated perforin expression in naïve CD8 T cells. The inhibition significance increases with hyperfunction in AF compared to MF or H (ANOVA interaction p=0.0890). D) Smo inhibition with cyclopamine during stimulation ablated perforin expression in effector CD8 T cells. The inhibition significance increases with hyperfunction in AF compared to MF or H (ANOVA interaction p=0.0314). All multiple comparisons are analyzed by 2-way ANOVA with Šídák’s post-test, \**p*<0.05, \*\**p*<0.01, \*\*\**p*<0.001.

## DISCUSSION

We report here that bulk circulating CD8 T cells from HCV-infected individuals exhibit differential gene expression patterns based on liver fibrosis severity after *in vitro* stimulation. Altered expression of genes associated with CD8 T cell function, survival, cellular metabolism, and cytoskeletal dynamics was identified through RNA-sequencing and subsequent bioinformatical discovery. Although our dataset is small, the gene expression differences were sufficiently significant to identify that gene expression patterns are not readily reverted after curative antiviral treatment, in-line with our previous observations of long-lasting hyperfunction^29^. This persistent dysfunction suggests that the observed CD8 T cell hyperfunction in HCV cirrhosis is due to cellular defects that are not readily reversed by the resolution of HCV viremia. Consequently, the persistence of dysfunctional CD8 T cells may contribute to remaining adverse clinical outcomes in HCV-infected individuals with cirrhosis, despite achieving SVR.

In chronic HCV infection, HCV-specific CD8 T cell responses are largely characterized by an exhaustive immune phenotype^10,38–40^. However, it is increasingly recognized that inflammatory cytokines facilitate T cell activation in the context of viral infection, enabling cells to circumvent antigen-dependency through an innate-like response^41^. Upregulated IFN-γ response genes in cells from patients with cirrhosis correlates with our previous finding of elevated proportions of IFN-γ^+^ cells in HCV infection^29^, although in a different context with this report showing an increase in genes downstream of IFN-γ signaling rather than IFN-γ genes themselves. Such parallels may be expected, as IFN-γ autocrine and paracrine signaling in mouse CD8 T cells has been shown to enhance motility and cytotoxicity, promoting T-bet and granzyme B expression^42,43^. Our data also identified an increased IFN-α responsiveness in CD8 T cells of HCV^+^ patients with cirrhosis compared to minimal fibrosis. It has been reported that IFN-α stimulation of PBMCs resulted in a phenotypic shift in CMV-/EBV-specific CD8 T cells from healthy individuals to that resembling cells in chronic HCV infection, with upregulated PD-1, Tim-3 and 2B4 expression^26^. IFN-α plays a role in supporting T cell receptor engagement and co-stimulation, similar to IL-12, by enhancing CD8 T cell differentiation and function^44^. IFN-α has also been shown to directly increase IFN-γ and granzyme production by naïve or antigen-experienced CD8 T cells^45,46^. Thus, these RNA-seq data suggests non-specific activation and function of CD8 T cells in HCV infection with advanced fibrosis.

Across several GSEA and GO referencing, changes in genes involved in cytoskeletal regulation further suggest dysregulation of cellular structure in CD8 T cells from HCV^+^ individuals with advanced fibrosis. These include identified upregulation in (e.g., leukocyte migration gene sets), as well as downregulation in four gene sets associated with the regulation and activation of actin nucleation (p<0.05, Fig. 2) and fifth as a trend towards the downregulation of genes associated with the positive regulation of actin polymerization. The ability of CD8 T cells to enact receptor-mediated intracellular signaling and release cytotoxic molecules hinges on tight actin filament regulation. Actin and microtubule reorganization, as well as inositol phosphate metabolism which were enriched in our RNA-seq data, play a central role in mediating the targeted synaptic release of cytolytic granules centrosomes^47^. Given these gene expression changes and the strong upregulation of Hh signaling in GSEA, which acts on the actin organization of CD8 T cells^36^, intracellular structural changes may be an important driver of CD8 T cell hyperfunction in advanced liver disease and impact their roles in responses to infection or cancer surveillance. Inositol triphosphate signaling, as well as phospholipase signaling, also play an important role in T cell receptor signaling in development and function^48^.

The degree of cellular heterogeneity in bulk circulating CD8 T cells, based on surface phenotypes alone, should be considered in the interpretation of such bulk RNA sequencing. In patients with multiple sclerosis, heterogeneity of gene expression has been documented in naïve CD4 T cells that suggest bias in gene expression potential can exist prior to antigen encounter^49^. Even cytotoxic CD8 T cells exhibit inherent heterogeneity, with IFN-γ-expressing cells being most responsible for cytolytic activities with the production of TNF-α, granzymes and perforin and chemokines associated with antimicrobial activity (e.g. CCL5), whereas IL-2-producing cells modulate immune response with IL-4, -3 and -11 cytokine production^50^. The increased expression of CCL5 suggesting potential co-expression with lytic molecules such as perforin is in agreement with our previous finding that CD8 T cells from cirrhotic patients express elevated levels of perforin^29^. Continuous systemic stimuli in chronic infection, such as persistently high levels of serum Hh ligands, could also influence cellular heterogeneity as well^51–53^. Additional studies are required to identify the CD8 T cell subsets responsible for generalized immune cell hyperfunction in cirrhosis, complemented with cell-discriminatory approaches such as single-cell RNA-sequencing should gene expression be of interest.

Lasting generalized CD8 T cell dysfunction in cirrhosis may reduce immunocompetence and lead to increased risk of community-acquired infections such as pneumonia^54,55^ and poor responses to routine vaccinations in HCV-infected individuals^56–60^. Landmark studies in murine chronic LCMV infection show how CD8 T cell dysfunction associates with several weakened aspects of the immune response^61,62^. HCV vaccines currently under development seek to emulate T cell responses required for protection against HCV, including reinfection^63^. However, it is thought that lasting CD8 T cell dysfunction after HCV cure will result in a failure to generate effective HCV vaccine responses in vulnerable populations^64^ and potentially contribute to the development of HCC and poor responses to immunotherapy therein.

To date, this is the first study to have probed generalized CD8 T cell gene expression patterns in chronic HCV infection in the context of liver disease severity. Evaluating the immune function in the context of chronic HCV-derived liver disease severity can be difficult due to confounding inter-individual immunological factors, such as minor infections/inflammation, and in the case of this study, small sample sizes. This has been reflected in many study data sets underrepresenting cirrhotics. Furthermore, the ability to conduct longitudinal studies to evaluate responses before and long after DAA therapy is difficult given the loss-to-follow-up phenomenon in this field, as seen in recent works^65^. It was also noted, although not statistically significant, that the gene expression patterns of cells from female patients all clustered closely in PCA, except for patient 136. A more in-depth analysis of possible sex-effects of fibrosis severity on CD8 T cell gene expression may be warranted in the future.

Restoration of general CD8 T cell function after therapeutic resolution of chronic HCV could have beneficial effects on the function of antigen-specific cells and help remediate complications associated with chronic HCV and cirrhosis, including HCC and other immune dysfunction. Identification of the underlying mechanisms of widespread CD8 T cell hyperactivation in chronic HCV infection with advanced liver fibrosis may provide insight on targets for immune restoration following antiviral therapy. Additionally, targets of immune restoration identified in the context of chronic HCV may inform potential restoration approaches across other liver disease etiologies, including HBV or HIV co-infections and MAFLD.

## ACKNOWLEDGMENTS

This study was funded by a Canadian Network on Hepatitis C (CanHepC) Pilot Grant (NHC-142832) and a Canadian Institutes of Health Research (CIHR) Operating Grant (PJT-419982). JL is funded by a CanHepC Graduate Scholarship, and a Queen Elizabeth II Graduate Scholarship in Science and Technology from the Government of Ontario. We sincerely appreciate the technical assistance of Sean Delaney (William L Stanford lab) in the RNA isolation and spike-in procedures for RNA-seq. We also sincerely appreciate the assistance of Christopher J Porter (Ottawa Bioinformatics Core Facility, Ottawa Hospital Research Institute (OHRI) and University of Ottawa) in the bioinformatical processing and analysis of RNA-seq data. We gratefully acknowledge the RNA-sequencing services provided the Donnelly Sequencing Centre (Toronto, Canada). We also gratefully acknowledge the RNA capillary electrophoresis services provided by StemCore Laboratories (RRID: SCR_012601) at the OHRI, and the services provided by the Flow Cytometry and Virometry Core Facility (RRID: SCR_023306) at the University of Ottawa for access to high-throughput multi-parameter flow cytometry tools and equipment.

## AUTHOR CONTRIBUTIONS

AV performed the RNA-seq experiments, trained staff and assisted in supervising experiments, and aided in the editing of the manuscript. JL performed RNA-seq analysis, RT-qPCR experiments, CD8 T cell culture experiments, analyzed data, and wrote and edited the manuscript. DR assisted in CD8 T cell culture experiments. WLS aided in the RNA-seq analysis and aided in the editing of the manuscript. CLC contributed to the conception of the study, led participant selection, recruitment, and specimen collection, and aided in the editing of the manuscript. AMC designed the project, supervised experiments, assisted with data analysis, and wrote and edited the manuscript.

## METHODS

### Study subjects

Study subjects (Tables 1) were treatment-naïve, chronically infected with HCV (>6 months HCV RNA^+^). All DAA-treated individuals studied achieved SVR unless otherwise specified. This research was conducted in accordance with the guidelines established by the Ottawa Health Science Network Research Ethics Board. Study participants were consented, and blood samples were collected by staff at The Ottawa Hospital Clinical Investigations Unit.

### CD8 T cell isolation and culture

PBMCs of study participants were isolated by Lymphoprep™ density gradient centrifugation (StemCell™ Technologies #07851, Canada), and cryopreserved in heat-inactivated fetal-bovine serum (HI-FBS, ThermoFisher Gibco™ #12484028, USA) + 10% (v/v) dimethyl sulfoxide (Sigma-Aldrich #472301, USA) at 1×10^7^ viable cells/ml. At the time of use, cryopreserved PBMCs were thawed and rested for 16h in RPMI 1640 (ThermoFisher Gibco™ #11875093) + 10% (v/v) HI-FBS + 100 U/ml penicillin-streptomycin (pen-strep, ThermoFisher Gibco™ #15140122) at 37°C, 5% CO_2_. CD8 T cells were then isolated by magnetic bead positive selection (StemCell™ Technologies #17853). Isolated CD8 T cells were cultured at 1×10^6^ cells/ml in complete RPMI (RPMI 1640 + 20% (v/v) HI-FBS + 100 U/ml pen-strep) at 37°C, 5% CO_2_ for all experiments.

### RNA sequencing of CD8 T cells

Isolated CD8 T cells were stimulated in culture with 0.5 μg/ml phytohemaglutanin-L (PHA-L, Sigma-Aldrich #L2769) for 18h. Following stimulation, total RNA was isolated using TRIzol™ Reagent (ThermoFisher Invitrogen™ #15596026) following manufacturer’s protocol. Total RNA yields were determined by spectrophotometer (ThermoFisher NanoDrop™ ND-1000) analysis of the A260/A280 ratio. To enable performance quality assessment during sequencing, spike-in control RNA (ThermoFisher Ambion™ #4456740) was added to all samples at a 1:100 dilution following manufacturer’s protocols.

RNA-sequencing was carried out by the Donnelly Sequencing Centre (DSC) in Toronto, Canada. Briefly, the purity and integrity (threshold RNA Integrity Number >8) of isolated total RNA was determined by microfluidic spectrophotometry on the 2100 Bioanalyzer (Agilent Technologies, USA). RNA-seq libraries were generated via mRNA isolation from total RNA by polyA-positive selection using the TruSeq™ RNA/DNA Library Preparation Kit (Illumina #RS-122-2001, USA) prior to sequencing using the NextSeq™ 550 system (Illumina).

### RNA-seq data analysis

Bioinformatical analysis of RNA-seq data was carried out in collaboration with the Ottawa Bioinformatics Core Facility (OHRI and University of Ottawa, Ottawa, Canada) using the R programming language. Read mapping was performed using the *Salmon* tool^66^ and quality control was subsequently performed using the *HISAT2* tool^67^. Fold-change analysis was then performed using the *DESeq2* tool^68^, applying for each gene a cutoff of ≥5 detectable reads in ≥2 samples for retention, which removes non-expressed and non-detectable, based on the *tximport* transcripts library^69^. Hierarchical clustering was calculated using all detectable transcripts. The top 500 most variably expressed genes across the samples was used to generate the principal component analysis plot. Statistically significant differentially expressed genes between study groups were identified using the *aleglm* method^70^ prior to Gene Set Enrichment Analysis^71,72^ and Gene Ontology enrichment analysis^73–75^.

### RT-qPCR of Hedgehog signaling genes in CD8 T cells

Isolated CD8 T cells were stimulated in culture with anti-CD3/CD28 antibodies. Briefly, high-binding 96-well plates were coated with 5 μg/ml anti-CD3 (Clone UCHT1, BD Pharmingen™ #555329, USA) in PBS (ThermoFisher Gibco™ #21600044) for 1h at 37°C, 5% CO_2_ prior to seeding cells for culture (conditions described above). Soluble anti-CD28 (Clone CD28.2, BD Pharmingen™ #555725) was then added to the cells at 2 μg/ml, and cells were cultured for 16h. Following stimulation, total RNA was isolated using the RNeasy Plus Micro kit (QIEGEN #74034, Netherlands) following manufacturer’s protocol. Total RNA yields were determined by spectrophotometry using NanoDrop™ One (ThermoFisher) analysis of the A260/A280 ratio, and RNA purity and integrity was determined by capillary electrophoresis on the 5200 Fragment Analyzer (Agilent Technologies) at StemCore Laboratories (OHRI). cDNA was generated using the iScript™ cDNA Synthesis Kit (Bio-Rad #1708890, USA) following manufacturer’s protocol, and gene expression was assessed by qPCR using the SYBR Green reporter (Bio-Rad, #1725270) on the CFX Connect (Bio-Rad) using the following PrimePCR™ (Bio-Rad) primer assays (Gene, Assay ID): PTCH1, qHsaCED0001809; GLI1, qHsaCID0011958; TBP (housekeeping gene), qHsaCID0007122. Fold-change of gene expression between study groups was calculated using the 2^-ΔCt^ method normalized to TBP.

### Inhibition of Hedgehog signaling in CD8 T cells

Isolated CD8 T cells were stimulated in culture with anti-CD3/CD28 antibodies as described above for 48h, with or without 50 μM cyclopamine (StemCell™ Technologies #72072) or Vismodegib (Selleck Chemicals #S1082, USA). The expression of IFN-γ and perforin in CD8 T cell subsets were analyzed by spectral flow cytometry on the Aurora (Cytek Bioscience, USA) using the following markers (Clone, Cat. #, BioLegend, USA): Zombie Aqua™ fixable viability dye (#423101), BV785 CD8 (RPA-T8, #301046), APC-Cy7 CCR7 (G043H7, #353212), BV650 CD45RA (HI100, #304136), IFN-γ (4S.B3, #502528), Perforin (B-D48, #353312). Data was analyzed with FlowJo™ v.10 software (BD) and statistical analysis was performed with Prism v.10 software (Dotmatics GraphPad™, USA).

## REFERENCS

1. Alberti, A. & Piovesan, S. Increased incidence of liver cancer after successful DAA treatment of chronic hepatitis C: Fact or fiction? Liver Int. Off. J. Int. Assoc. Study Liver 37, 802–808 (2017).

2. Aleman, S. et al. A risk for hepatocellular carcinoma persists long-term after sustained virologic response in patients with hepatitis C-associated liver cirrhosis. Clin. Infect. Dis. Off. Publ. Infect. Dis. Soc. Am. 57, 230–236 (2013).

3. Romano, A. et al. Newly diagnosed hepatocellular carcinoma in patients with advanced hepatitis C treated with DAAs: A prospective population study. J. Hepatol. 69, 345–352 (2018).

4. Tokita, H. et al. Risk factors for the development of hepatocellular carcinoma among patients with chronic hepatitis C who achieved a sustained virological response to interferon therapy. J. Gastroenterol. Hepatol. 20, 752–758 (2005).

5. Reig, M., Boix, L. & Bruix, J. The impact of direct antiviral agents on the development and recurrence of hepatocellular carcinoma. Liver Int. Off. J. Int. Assoc. Study Liver 37 Suppl 1, 136–139 (2017).

6. Sipeki, N., Antal-Szalmas, P., Lakatos, P. L. & Papp, M. Immune dysfunction in cirrhosis. World J. Gastroenterol. 20, 2564–2577 (2014).

7. Bernsmeier, C. et al. CD14+ CD15-HLA-DR-myeloid-derived suppressor cells impair antimicrobial responses in patients with acute-on-chronic liver failure. Gut 67, 1155–1167 (2018).

8. Rolas, L. et al. NADPH oxidase depletion in neutrophils from patients with cirrhosis and restoration via toll-like receptor 7/8 activation. Gut 67, 1505–1516 (2018).

9. Vergis, N. et al. Defective monocyte oxidative burst predicts infection in alcoholic hepatitis and is associated with reduced expression of NADPH oxidase. Gut 66, 519–529 (2017).

10. Cox, A. L. et al. Comprehensive analyses of CD8+ T cell responses during longitudinal study of acute human hepatitis C. Hepatol. Baltim. Md 42, 104–112 (2005).

11. Gruener, N. H. et al. Sustained dysfunction of antiviral CD8+ T lymphocytes after infection with hepatitis C virus. J. Virol. 75, 5550–5558 (2001).

12. Penna, A. et al. Dysfunction and functional restoration of HCV-specific CD8 responses in chronic hepatitis C virus infection. Hepatol. Baltim. Md 45, 588–601 (2007).

13. Rehermann, B. Chronic infections with hepatotropic viruses: mechanisms of impairment of cellular immune responses. Semin. Liver Dis. 27, 152–160 (2007).

14. Shen, T., Chen, X., Xu, Q., Lu, F. & Liu, S. Distributional characteristics of CD25 and CD127 on CD4+ T cell subsets in chronic HCV infection. Arch. Virol. 155, 627–634 (2010).

15. Wedemeyer, H. et al. Impaired effector function of hepatitis C virus-specific CD8+ T cells in chronic hepatitis C virus infection. J. Immunol. Baltim. Md 1950 169, 3447–3458 (2002).

16. Cooper, S. et al. Analysis of a successful immune response against hepatitis C virus. Immunity 10, 439–449 (1999).

17. Thimme, R. et al. Determinants of viral clearance and persistence during acute hepatitis C virus infection. J. Exp. Med. 194, 1395–1406 (2001).

18. Abdel-Hakeem, M. S., Bédard, N., Murphy, D., Bruneau, J. & Shoukry, N. H. Signatures of protective memory immune responses during hepatitis C virus reinfection. Gastroenterology 147, 870–881.e8 (2014).

19. Abdel-Hakeem, M. S. et al. Comparison of immune restoration in early versus late alpha interferon therapy against hepatitis C virus. J. Virol. 84, 10429–10435 (2010).

20. Hengst, J. et al. Nonreversible MAIT cell-dysfunction in chronic hepatitis C virus infection despite successful interferon-free therapy. Eur. J. Immunol. 46, 2204–2210 (2016).

21. Spaan, M. et al. Immunological Analysis During Interferon-Free Therapy for Chronic Hepatitis C Virus Infection Reveals Modulation of the Natural Killer Cell Compartment. J. Infect. Dis. 213, 216–223 (2016).

22. Lucas, M. et al. Pervasive influence of hepatitis C virus on the phenotype of antiviral CD8+ T cells. J. Immunol. Baltim. Md 1950 172, 1744–1753 (2004).

23. Rehermann, B. et al. Quantitative analysis of the peripheral blood cytotoxic T lymphocyte response in patients with chronic hepatitis C virus infection. J. Clin. Invest. 98, 1432–1440 (1996).

24. Sumida, K. et al. Characteristics of splenic CD8+ T cell exhaustion in patients with hepatitis C. Clin. Exp. Immunol. 174, 172–178 (2013).

25. Zhao, B.-B. et al. T lymphocytes from chronic HCV-infected patients are primed for activation-induced apoptosis and express unique pro-apoptotic gene signature. PloS One 8, e77008 (2013).

26. Owusu Sekyere, S. et al. Type I Interferon Elevates Co-Regulatory Receptor Expression on CMV- and EBV-Specific CD8 T Cells in Chronic Hepatitis C. Front. Immunol. 6, 270 (2015).

27. Moreno-Cubero, E. et al. According to Hepatitis C Virus (HCV) Infection Stage, Interleukin-7 Plus 4-1BB Triggering Alone or Combined with PD-1 Blockade Increases TRAF1low HCV-Specific CD8+ Cell Reactivity. J. Virol. 92, e01443–17 (2018).

28. Burke Schinkel, S. C., Carrasco-Medina, L., Cooper, C. L. & Crawley, A. M. Generalized Liver- and Blood-Derived CD8+ T-Cell Impairment in Response to Cytokines in Chronic Hepatitis C Virus Infection. PloS One 11, e0157055 (2016).

29. Vranjkovic, A. et al. Direct-Acting Antiviral Treatment of HCV Infection Does Not Resolve the Dysfunction of Circulating CD8+ T-Cells in Advanced Liver Disease. Front. Immunol. 10, 1926 (2019).

30. Doyle, M.-A., Galanakis, C., Mulvihill, E., Crawley, A. & Cooper, C. L. Hepatitis C Direct Acting Antivirals and Ribavirin Modify Lipid but not Glucose Parameters. Cells 8, E252 (2019).

31. Carballo, G. B., Honorato, J. R., de Lopes, G. P. F. & Spohr, T. C. L. de S. E. A highlight on Sonic hedgehog pathway. Cell Commun. Signal. CCS 16, 11 (2018).

32. Briscoe, J. & Thérond, P. P. The mechanisms of Hedgehog signalling and its roles in development and disease. Nat. Rev. Mol. Cell Biol. 14, 416–429 (2013).

33. Teperino, R., Aberger, F., Esterbauer, H., Riobo, N. & Pospisilik, J. A. Canonical and non-canonical Hedgehog signalling and the control of metabolism. Semin. Cell Dev. Biol. 33, 81–92 (2014).

34. Crompton, T., Outram, S. V. & Hager-Theodorides, A. L. Sonic hedgehog signalling in T-cell development and activation. Nat. Rev. Immunol. 7, 726–735 (2007).

35. Mengrelis, K. et al. Sonic Hedgehog Is a Determinant of γδ T-Cell Differentiation in the Thymus. Front. Immunol. 10, 1629 (2019).

36. de la Roche, M. et al. Hedgehog signaling controls T cell killing at the immunological synapse.Science 342, 1247–1250 (2013).

37. Martelli, A. M. et al. Understanding the Roles of the Hedgehog Signaling Pathway during T-Cell Lymphopoiesis and in T-Cell Acute Lymphoblastic Leukemia (T-ALL). Int. J. Mol. Sci. 24, 2962 (2023).

38. Radziewicz, H. et al. Liver-infiltrating lymphocytes in chronic human hepatitis C virus infection display an exhausted phenotype with high levels of PD-1 and low levels of CD127 expression. J. Virol. 81, 2545–2553 (2007).

39. Hensel, N. et al. Memory-like HCV-specific CD8+ T cells retain a molecular scar after cure of chronic HCV infection. Nat. Immunol. 22, 229–239 (2021).

40. Reid, A. & Humblin, E. Exhausted T cells never fully recover. Nat. Rev. Immunol. 21, 408 (2021).

41. Berg, R. E. & Forman, J. The role of CD8 T cells in innate immunity and in antigen non-specific protection. Curr. Opin. Immunol. 18, 338–343 (2006).

42. Bhat, P., Leggatt, G., Waterhouse, N. & Frazer, I. H. Interferon-γ derived from cytotoxic lymphocytes directly enhances their motility and cytotoxicity. Cell Death Dis. 8, e2836–e2836 (2017).

43. Curtsinger, J. M., Agarwal, P., Lins, D. C. & Mescher, M. F. Autocrine IFN-γ Promotes Naive CD8 T Cell Differentiation and Synergizes with IFN-α To Stimulate Strong Function. J. Immunol. 189, 659–668 (2012).

44. Huber, J. P. & Farrar, J. D. Regulation of effector and memory T-cell functions by type I interferon. Immunology 132, 466–474 (2011).

45. Hervas-Stubbs, S. et al. Effects of IFN-α as a signal-3 cytokine on human naïve and antigen-experienced CD8(+) T cells. Eur. J. Immunol. 40, 3389–3402 (2010).

46. Kohlmeier, J. E., Cookenham, T., Roberts, A. D., Miller, S. C. & Woodland, D. L. Type I interferons regulate cytolytic activity of memory CD8(+) T cells in the lung airways during respiratory virus challenge. Immunity 33, 96–105 (2010).

47. de la Roche, M., Asano, Y. & Griffiths, G. M. Origins of the cytolytic synapse. Nat. Rev. Immunol.16, 421–432 (2016).

48. Huang, Y. H. & Sauer, K. Lipid signaling in T-cell development and function. Cold Spring Harb. Perspect. Biol. 2, a002428 (2010).

49. Sood, A. et al. Differential interferon-gamma production potential among naïve CD4+ T cells exists prior to antigen encounter. Immunol. Cell Biol. 97, 931–940 (2019).

50. Nicolet, B. P. et al. CD29 identifies IFN-γ-producing human CD8+ T cells with an increased cytotoxic potential. Proc. Natl. Acad. Sci. U. S. A. 117, 6686–6696 (2020).

51. Navas, M.-C. et al. Hepatitis C Virus Infection and Cholangiocarcinoma: An Insight into Epidemiologic Evidences and Hypothetical Mechanisms of Oncogenesis. Am. J. Pathol. 189, 1122–1132 (2019).

52. Omenetti, A. & Diehl, A. M. The adventures of sonic hedgehog in development and repair. II. Sonic hedgehog and liver development, inflammation, and cancer. Am. J. Physiol. Gastrointest. Liver Physiol. 294, G595–598 (2008).

53. Pereira, T. de A. et al. Viral factors induce Hedgehog pathway activation in humans with viral hepatitis, cirrhosis, and hepatocellular carcinoma. Lab. Investig. J. Tech. Methods Pathol. 90, 1690–1703 (2010).

54. Bonnel, A. R., Bunchorntavakul, C. & Reddy, K. R. Immune dysfunction and infections in patients with cirrhosis. Clin. Gastroenterol. Hepatol. Off. Clin. Pract. J. Am. Gastroenterol. Assoc. 9, 727–738 (2011).

55. Borzio, M. et al. Bacterial infection in patients with advanced cirrhosis: a multicentre prospective study. Dig. Liver Dis. Off. J. Ital. Soc. Gastroenterol. Ital. Assoc. Study Liver 33, 41–48 (2001).

56. Cheong, H.-J. et al. Humoral and cellular immune responses to influenza vaccine in patients with advanced cirrhosis. Vaccine 24, 2417–2422 (2006).

57. De Maria, N., Idilman, R., Colantoni, A., Harig, J. M. & Van Thiel, D. H. Antibody response to hepatitis B virus vaccination in individuals with hepatitis C virus infection. Hepatol. Baltim. Md 32, 444–445 (2000).

58. Greenbaum, E. et al. Severe influenza infection in a chronic hepatitis C carrier: failure of protective serum HI antibodies after IM vaccination. J. Clin. Virol. Off. Publ. Pan Am. Soc. Clin. Virol. 29, 23–26 (2004).

59. Moorman, J. P. et al. Impaired hepatitis B vaccine responses during chronic hepatitis C infection: involvement of the PD-1 pathway in regulating CD4(+) T cell responses. Vaccine 29, 3169–3176 (2011).

60. Wiedmann, M. et al. Decreased immunogenicity of recombinant hepatitis B vaccine in chronic hepatitis C. Hepatol. Baltim. Md 31, 230–234 (2000).

61. Fallet, B. et al. Interferon-driven deletion of antiviral B cells at the onset of chronic infection. Sci. Immunol. 1, eaah6817 (2016).

62. Moseman, E. A., Wu, T., de la Torre, J. C., Schwartzberg, P. L. & McGavern, D. B. Type I interferon suppresses virus-specific B cell responses by modulating CD8+ T cell differentiation. Sci. Immunol. 1, eaah3565 (2016).

63. Shoukry, N. H. et al. Memory CD8+ T cells are required for protection from persistent hepatitis C virus infection. J. Exp. Med. 197, 1645–1655 (2003).

64. Houghton, M. Prospects for prophylactic and therapeutic vaccines against the hepatitis C viruses.Immunol. Rev. 239, 99–108 (2011).

65. Rosenberg, B. R. et al. Longitudinal transcriptomic characterization of the immune response to acute hepatitis C virus infection in patients with spontaneous viral clearance. PLoS Pathog. 14, e1007290 (2018).

66. Patro, R., Duggal, G., Love, M. I., Irizarry, R. A. & Kingsford, C. Salmon provides fast and bias-aware quantification of transcript expression. Nat. Methods 14, 417–419 (2017).

67. Kim, D., Paggi, J. M., Park, C., Bennett, C. & Salzberg, S. L. Graph-based genome alignment and genotyping with HISAT2 and HISAT-genotype. Nat. Biotechnol. 37, 907–915 (2019).

68. Love, M. I., Huber, W. & Anders, S. Moderated estimation of fold change and dispersion for RNA-seq data with DESeq2. Genome Biol. 15, 550 (2014).

69. Soneson, C., Love, M. I. & Robinson, M. D. Differential analyses for RNA-seq: transcript-level estimates improve gene-level inferences. F1000Research 4, 1521 (2015).

70. Zhu, A., Ibrahim, J. G. & Love, M. I. Heavy-tailed prior distributions for sequence count data: removing the noise and preserving large differences. Bioinforma. Oxf. Engl. 35, 2084–2092 (2019).

71. Subramanian, A. et al. Gene set enrichment analysis: a knowledge-based approach for interpreting genome-wide expression profiles. Proc. Natl. Acad. Sci. U. S. A. 102, 15545–15550 (2005).

72. Mootha, V. K. et al. PGC-1alpha-responsive genes involved in oxidative phosphorylation are coordinately downregulated in human diabetes. Nat. Genet. 34, 267–273 (2003).

73. Ashburner, M. et al. Gene ontology: tool for the unification of biology. The Gene Ontology Consortium. Nat. Genet. 25, 25–29 (2000).

74. Gene Ontology Consortium et al. The Gene Ontology knowledgebase in 2023. Genetics 224, iyad031 (2023).

75. Thomas, P. D. et al. PANTHER: Making genome-scale phylogenetics accessible to all. Protein Sci. Publ. Protein Soc. 31, 8–22 (2022).

