## Supplemental figures for "Differential and lasting gene expression changes in circulating CD8 T cells in chronic HCV infection with cirrhosis and related insights on the role of Hedgehog signaling"

### Figure S1:

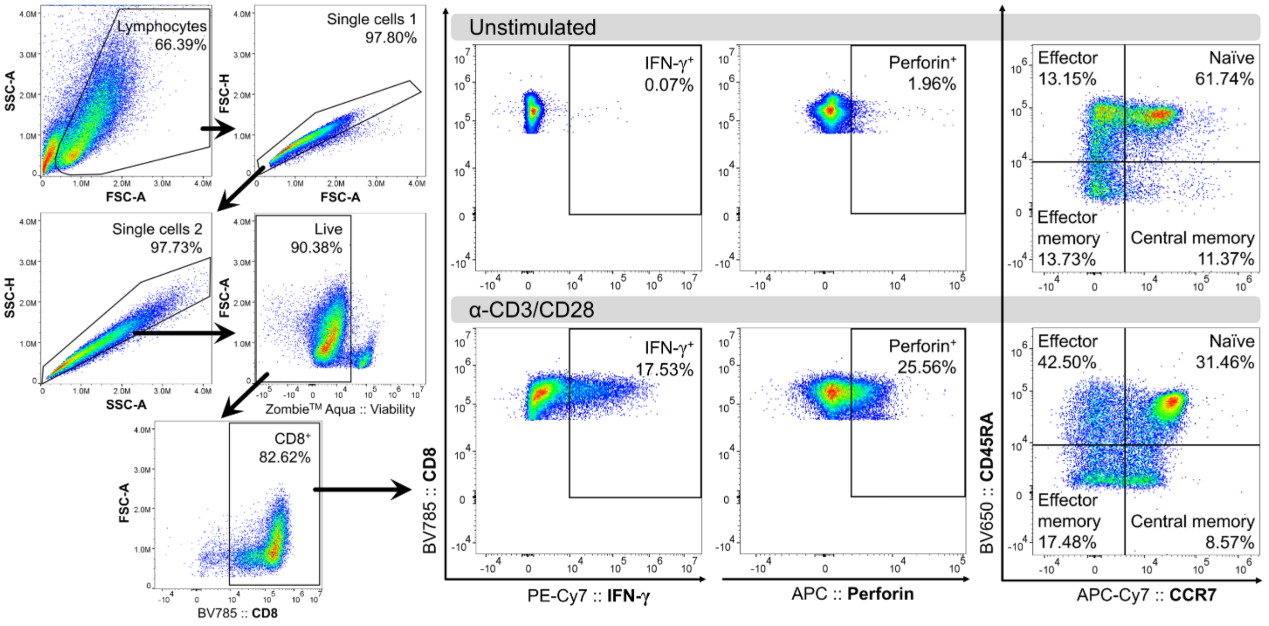
